# Would conserving natural land cover in landscapes conserve biodiversity?

**DOI:** 10.1101/436410

**Authors:** Rafael X. De Camargo, David J. Currie

## Abstract

It is generally accepted that protecting natural land cover would protect biodiversity. This would only be true as a general statement if the relationship between richness and natural land cover were *monotonic positive* and *scale*- and *method-independent*. Assertions about habitat loss causing species losses often come from broad-scale assessment of richness (e.g., from range maps) combined with patterns of natural habitat conversion. Yet, the evidence about species loss following habitat loss or fragmentation typically comes from fine-scale experiments. Here, we test whether broad-extent relationships between avian species richness and natural land cover are independent of: 1) whether species distribution data come from systematic censuses (atlases) versus range maps, and 2) the grain size of the analysis. We regressed census-based and range map-based avian species richness against the proportion of natural land cover and temperature. Censused richness at the landscape level was obtained from Breeding Bird Atlases of Ontario and New York State. Range-map richness derived from BirdLife International range maps. Comparisons were made across different spatial grains: 25-km^2^, 100-km^2^, and 900-km^2^. Over regional extents, range-map richness relates strongly to temperature, irrespective of spatial grain. Censused species richness relates to temperature less strongly. Range-map richness is a *negative* function of the proportion of natural land cover, while realized richness is a peaked function. The two measures of richness are not monotonically related to each other. In conclusion, the data do not indicate that, in practice, landscapes with greater natural land cover in southern Ontario or in New York State have higher species richness. Moreover, different data types can lead to dramatically different relationships between richness and natural land cover. We argue that the argument that habitat loss is the main driver of species loss has become a panchreston. It may be misguiding conservation biology strategies by focusing on a threat that is too general to be usefully predictive.

## INTRODUCTION

Wilcove et al. [1] wrote: “habitat loss is the single greatest threat to biodiversity…”. This hypothesis seems to have been accepted as a truism. Since 1990s habitat loss has been cited as “the major threat” [1,2], “ the main cause” [3,4], or “the principal driver” of biodiversity loss [5–7], to mention just a few examples.

A corollary of the hypothesis that habitat loss is the main cause of species loss is that conserving natural land cover (a first approximation of habitat) will conserve biodiversity. If that is true, a critical prediction can then be tested: in landscapes in which human activities have removed part of the natural habitat, areas with more natural land cover should have higher species richness. At least three types of data can be brought to bear on this prediction. The first type is experimentally manipulated habitat patches of varying areas (usually at fairly small scales). Patch-level studies (1-50ha) consistently find that larger patches of uninterrupted habitat have more species than smaller patches [8–11]. However, conservation planning does not generally focus on individual patches of habitat; rather, planning more often involves landscapes (areas on the order of 10–1,000 km^2^) with mixtures of land covers [12], consequently patch-scale is not the focus in this study. A second category of data involves systematic field observations of species presence/absence in replicated landscapes (typically, 10^1^-10^3^ km^2^), which we shall call *censused richness*. Examples include regional or national breeding bird atlases, which may be called realised richness since it represents observed local assemblages of species. Finally, there are also stacked species range maps, typically over broad extents (e.g., continental), which we shall call *range-map richness*. Since, broad-scale range maps (e.g., IUCN, BirdLife International) are typically resolved at fairly coarse grains, e.g. ~10^4^ km^2^ [13], this represents the potential richness that is regionally available to occupy a landscape.

Biodiversity loss and its causes are clearly grain size dependent. There are many examples of extirpations of individual species, or even entire assemblages, associated with a variety of anthropogenic causes [14–18]. In general, these extinctions are local. At local scales, there is no clear consensus on the environmental determinants and mechanisms giving rise to pattern in richness, or to patterns of species extinctions [19–22]; however, in a recent study, land-use and land-use intensity have been proposed as potential drivers of local assembly patterns worldwide [23]. Similarly, at the landscape level, questions like “how much habitat is enough?” [24] is still controversial [25,26], suggesting that, to a certain extent, land cover and suitable habitats should be important to species maintenance. On the other hand, at meso- or larger scales, species extirpations/extinctions have been relatively rare [27]. Abiotic variables may better explain the occurrence of individual species [28]. Contemporary climate (e.g., temperature and precipitation) [29–31] and historical/evolutionary drivers [32,33] are the main competing hypotheses proposed to explain variation in species richness at broad scales.

The literature suggests that the relationship between richness and natural land cover should be scale-independent. At fine scales, species-area relationship modelling (e.g., richness as a power function of forest cover) has consistently shown a large negative effect of forest loss on biodiversity [34–38]. In a given locality, it must be true that complete elimination of natural land cover leads to species’ extirpation. However, researchers have long claimed that deforestation leads to loss of biodiversity worldwide [39–44]. Early species-area estimates from tropical forest deforestation in 1980s predicted that Earth could lose up to 20% of its species by the year of 2000 [39]. Later calculations from logging and deforestation predicted losses of between 37 and 50% of tree species of the Brazilian Amazon [44]. Pimm & Raven [40] predicted 18% extinction by 2100 in tropical hotspots due to due to forest loss. All of these predictions used coarse-grain species’ ranges modelled as a function of the amount of forest cover. In a recent paper [45], using (coarse-grained) IUCN range maps predicted that 121–219 species will become threatened under current rates of forest loss over the next 30 years in Borneo, the central Amazon, and the Congo Basin.

In this study, we tested whether the relationship between avian species richness and natural land cover is dependent upon the data type: fine-grained distribution data, versus coarser-grain range map data, sampled in grid cells of varying size. The literature assumes that the relationship between avian species richness and natural land cover is the same using fine-grained (censused) species’ distributions and coarser-grained (potential) species’ ranges. Since continental-scale studies of richness from range maps have generally observed strong richness-climate relationships, one might expect range-map richness to be more strongly related to temperature than to land cover. In contrast, one might expect censused richness to be more strongly related to land cover. To test whether these relationships depend on grain size (cell sizes) at which species are recorded, we also tested the consistency of the richness-land cover relationship across grain sizes of 25-km^2^, 100-km^2^ and 900-km^2^. Finally, to test the consistency of our results, we compare fitted models from separate datasets covering southern Ontario and New York State.

## METHODS

### Study region and species richness

The study geographical region includes southern Ontario, Canada (200,000 km^2^), and New York State, USA (125,400 km^2^) (Fig 1). To calculate censused, landscape-level avian species richness, we used species distribution data from the Ontario Breeding Bird Atlas (OBBA, [46]) and the New York State Breeding Bird Atlas (NYBBA, [47]). Both atlases were based on systematic surveys conducted between 2000 and 2005.

**Fig 1.**
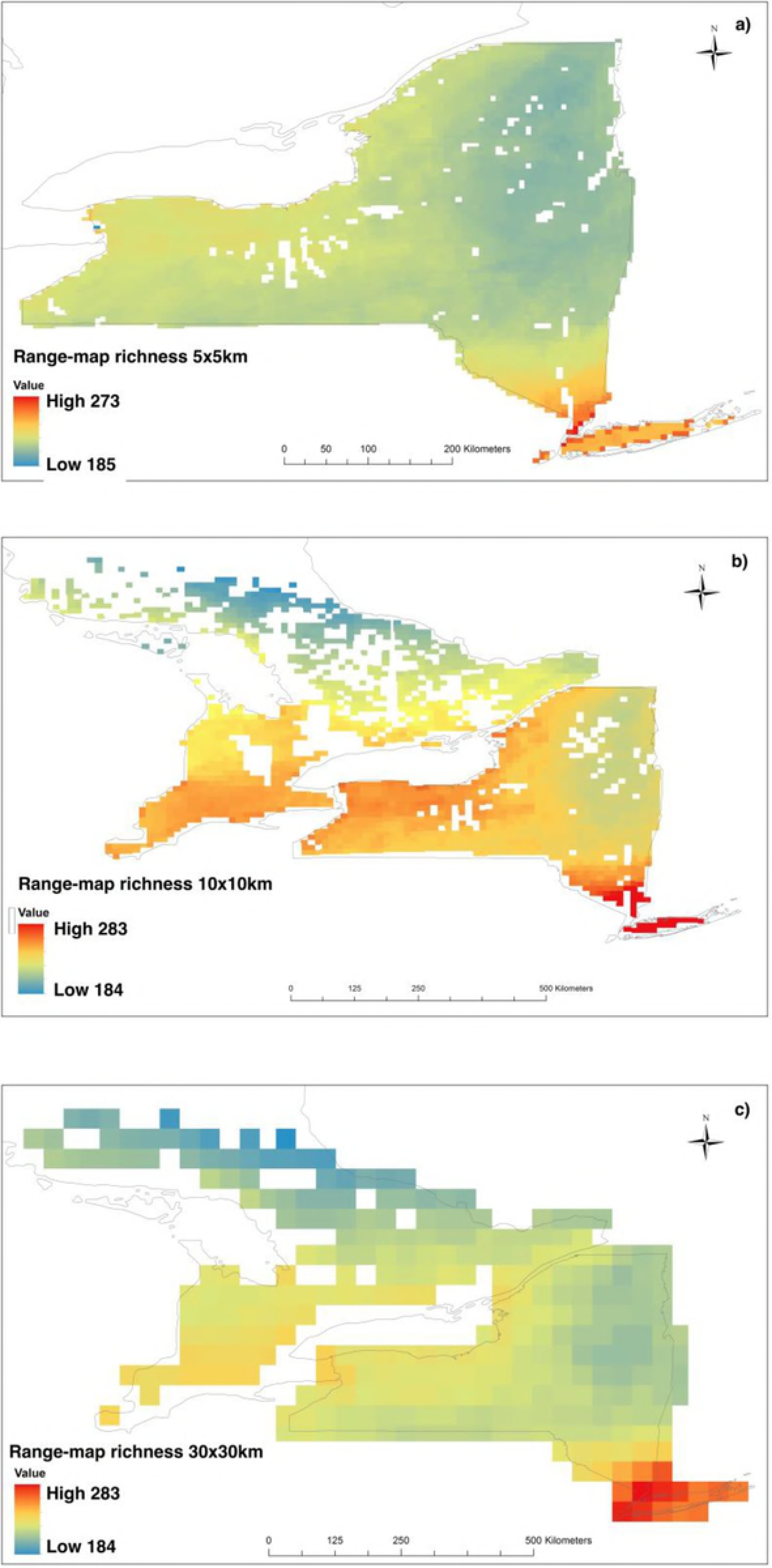
The proportion of natural land cover covering the study area according to the global 1-km consensus land cover data set [48]. The projection is WGS84 datum.

The NYBBA sampled birds on a 5×5km grid; the OBBA used a cell size of 10×10km. Both atlases used experienced birders to identify the breeding bird species occurring within each quadrat. Since the sampling was designed to sample all habitats in a grid cell, and hopefully to find all species breeding there, we treat species not observed in a cell as being truly absent [49]. Richness in a quadrat represents the total number of species presences observed in that quadrat.

Sampling effort varies both within and between the two atlases. For the NYBBA, atlassers were assigned to survey one or more NYBBA quadrats and were expected to spend at least 8h in each block, visiting each habitat present, and recording at least 76 species. For the ABBO surveys, each volunteer was assigned to search a specific 100-km^2^quadrat as completely as possible for evidence of all species breeding therein. Volunteers were instructed to search in particular for regionally rare species. Any species that was observed in a given quadrat in 2000- 2005 was considered present. We excluded ABBO quadrats with <20 hours and > 600 hours of bird censusing effort (median effort ≅45 hours). Since the OBBA quadrats were four times larger than the NYBBA quadrats, the effort per unit area was similar in the two atlases.

In order to compare *censused richness* between atlases, we resampled the NYBBA at 10×10 km quadrat size (same cell-size as ABBO). We calculated censused richness by counting the number of unique species’ presences from the original survey quadrats within each new 100-km^2^ grid cells in New York State.

*Potential bird species richness* was extracted from species’ range maps in the BirdLife International World Bird Database (available online at http://www.birdlife.org/datazone). We overlaid species’ ranges on the 25-km^2^ (NYBBA) and 100-km^2^ (OBBA) quadrats. We also resampled New York using 10×10 km, and both Ontario and New York with 30×30km grid cells (900-km^2^). *Range-map richness* in a quadrat represents the total number of species’ ranges that overlap that quadrat.

### Richness predictors

To estimate natural land cover, we used a global consensus land cover data set [48] (Fig. 1). The dataset is composed of 12 land-cover classes, observed at a spatial resolution of 30 arc-seconds (~1-km^2^ pixels at the equator). The land-cover classes are: 1. Evergreen / Deciduous Needleleaf trees, 2. Evergreen broadleaf trees, 3. Deciduous broadleaf trees, 4. Mixedwood/other trees, 5. Shrubs, 6. Herbaceous vegetation, 7. Cultivated and Managed Vegetation, 8. Regularly flooded vegetation, 9. Urban/Built-up, 10. Snow/Ice, 11. Barren, and 12. Open Water. The proportion of pixels in each land cover class was determined within each quadrat (0-100%). To obtain the proportion of natural land cover in each of our sampling units, we summed the percentages of classes 1-6 and 8. We excluded grid cells containing more than 10% water. The total numbers of grid cells are: 4,822 for NY (5×5km), 985 and 1,075 for ON and NY, respectively, at 10×10km scale, and 251 covering ON and NY (30×30 km).

Temperature has been long known as a main correlate, perhaps the driver, of species richness patterns at broad-scales [28,30]. Hence, we used Mean Annual Temperature (MAT) from the WorldClim database (Fig 2, [50]), as a predictor of avian richness for each grid cell at different spatial scale.

**Fig 2.**
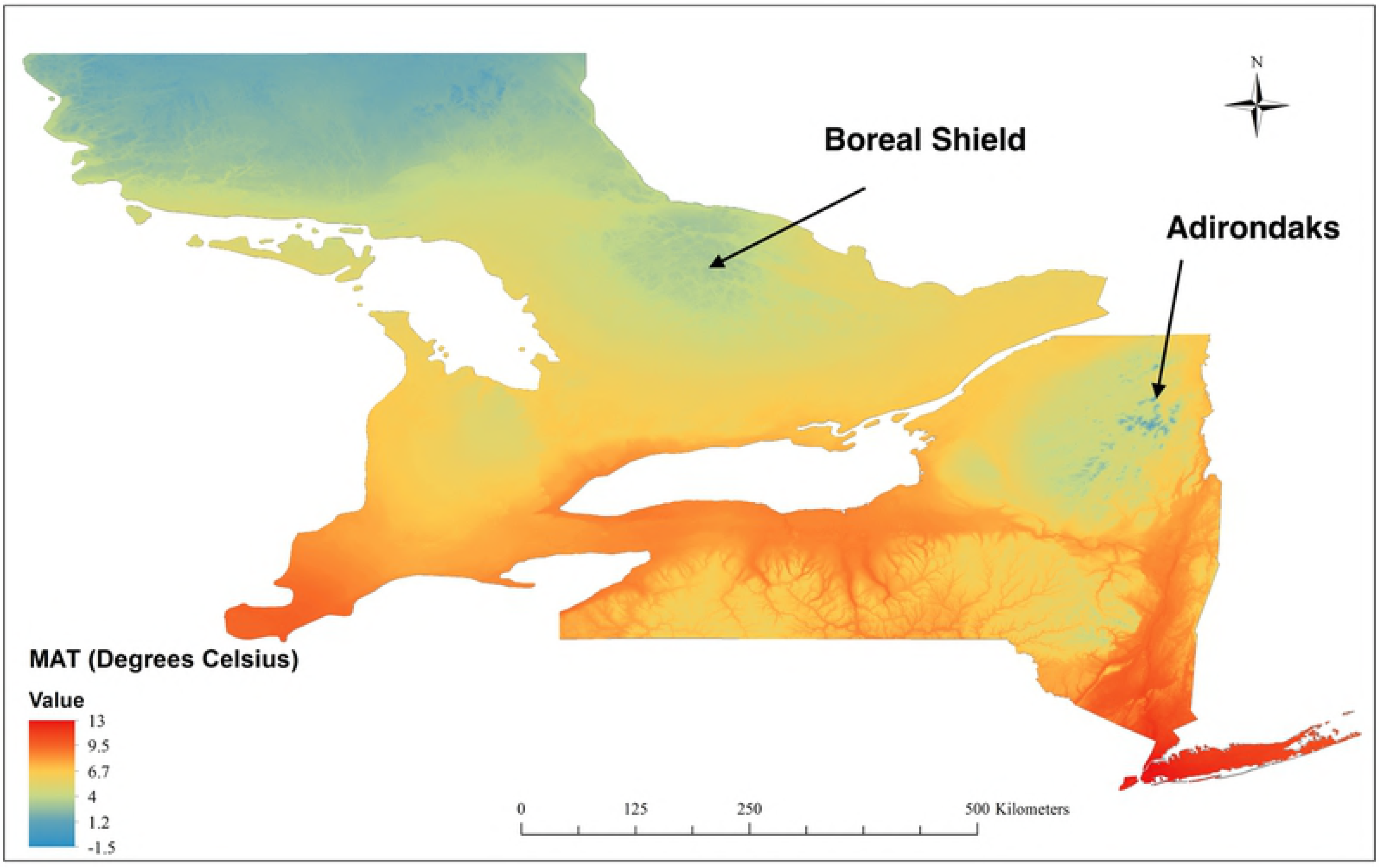
Mean Annual Temperature (MAT) covering the study area according to WorldClim [50]. The projection is WGS84 datum.

### Statistical Analysis

We used Ordinary Least Squares (OLS) regression models to relate censused and potential bird species richness to the environmental predictors: MAT and proportion of natural land cover. We fitted censused richness as a quadratic function of temperature and of the proportion of natural land cover, as these relationships are non-linear. We also fitted multiple regression models to determine the variance explained by the predictor variables together. Spatial data, including satellite images and climate raster files were treated in ArcGIS, and all stats were performed in R [51].

Spatial autocorrelation can affect models coefficients in spatial analyses [52]. Hence, we also corrected the richness models for spatial autocorrelation by fitting simultaneous autoregressive error models (SARerr) proposed by Kissling and Carl [53] in R (function “errorsarlm”).

## RESULTS

The relationship between avian species richness and natural land cover depends on the type of data from which richness gradients are generated. *Range-map richness* (Fig 3) clearly reflects the climatic gradients in the region (Fig 2) rather than land cover (Fig 1). Like climate, the spatial variation of range-map richness is strongly autocorrelated in space, across cell sizes from 25 km^2^ - 900 km^2^ (Fig 3a-c). Multiple regressions showed that range-map richness relates strongly to temperature, and it is *negatively* related to the proportion of natural land cover (Table 1, S3 Fig), contradicting the expected positive richness-land cover relationship. Moreover, most of the small amount of variance in range-map richness explained by natural land cover (Table 1) reflects collinearity between land cover and climate: forested areas in New York and Ontario are mainly in cold areas (S1 Fig). The observed patterns are very similar across 25-km^2^, 100-km^2^, and 900km^2^ cell sizes (Fig 3a-c, Table 1).

**Table 1.** OLS regression models between **range-map richness** as a function of temperature (MAT, top line at each grain size) and the proportion of natural land cover (LC, middle line at each grain), or both (bottom line in each grain) in grid cells of southern Ontario and New York State. Terms in parentheses are not statistically significant (p>0.05).

| Grain Size | Location | Standardized coefficients | | $r^2$ | AICc |
| --- | --- | --- | --- | --- | --- |
|  |  | MAT | LC |  |  |
| 5x5km | NY (n=4,822) | 4.58 |  | 0.55 | 31475 |
|  |  |  | -0.19 | 0.21 | 34177 |
|  |  | 4.41 | -0.02 | 0.55 | 31461 |
| 10x10km | NY (n=1,075) | 4.48 |  | 0.52 | 6978 |
|  |  |  | -0.21 | 0.25 | 7463 |
|  |  | 4.16 | -0.03 | 0.53 | 6971 |
|  | ON (n=985) | 7.72 |  | 0.77 | 6512 |
|  |  |  | -0.33 | 0.65 | 6932 |
|  |  | 6.14 | -0.08 | 0.78 | 6460 |
| 30x30km | NY (n=165) | 8.18 |  | 0.69 | 1192 |
|  |  |  | -0.36 | 0.35 | 1314 |
|  |  | 8.01 | (-0.01) | 0.69 | 1194 |
|  | ON (n=138) | 7.62 |  | 0.77 | 912 |
|  |  |  | -0.36 | 0.65 | 973 |
|  |  | 6.22 | -0.08 | 0.78 | 908 |

**Fig 3.**
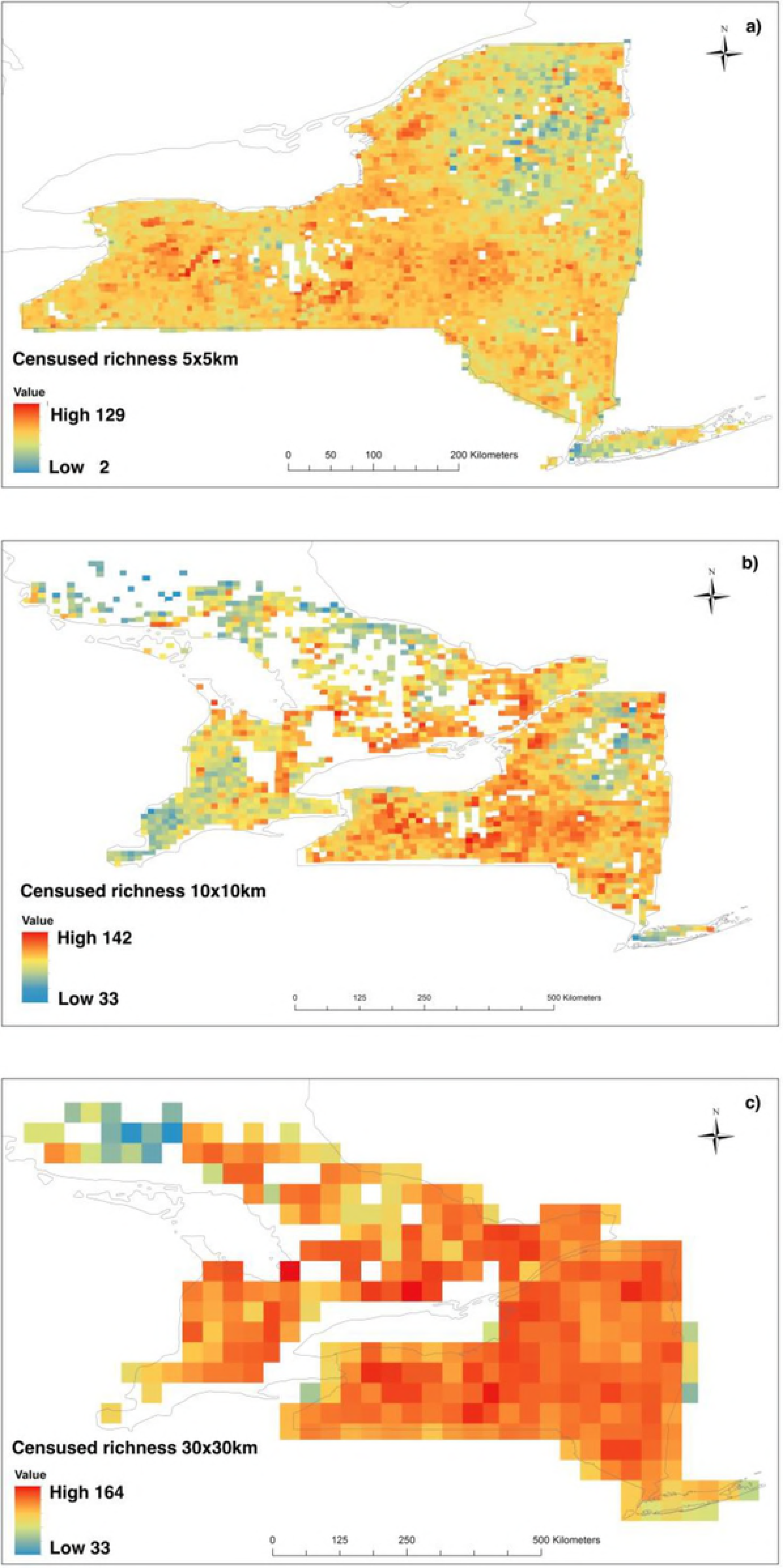
Distribution of potential avian species richness in a) 4,822 cells of 25-km^2^ in NY, b) 2,060 cells of 100-km^2^ in ON nad NY, and c) 303 cells of 900-km^2^ in ON and NY

The geographic variation of *censused richness* (obtained from fine-grained atlases) is less spatially structured (Fig 4), and it does reflect, to a certain degree, the spatial variation in the proportion of natural land cover in the study area (cf. Fig 1). Temperature still explains more variance in the models than the proportion of natural land cover (Table 2), but the censused richness-temperature relationships are peaked (Fig 5a,c,e). Natural land cover increases the r-squared of the models between by 4-13% (Table 2). Yet, high richness landscapes are interspersed throughout the study area, and they are not predominantly associated with the areas having a high proportion of natural land cover (e.g. the Adirondaks of north-eastern NY, the Catskills of south-eastern NY, and northern Ontario). The pattern is consistent across 25-km^2^, 100-km^2^, and 900km^2^ cell sizes (Fig 4a-c, Table 2). Moreover, the relationships between censused richness and land cover remained peaked even when the datasets were subset in order to encompass only the coldest or warmest regions of the study area (S2 Fig).

**Table 2.**
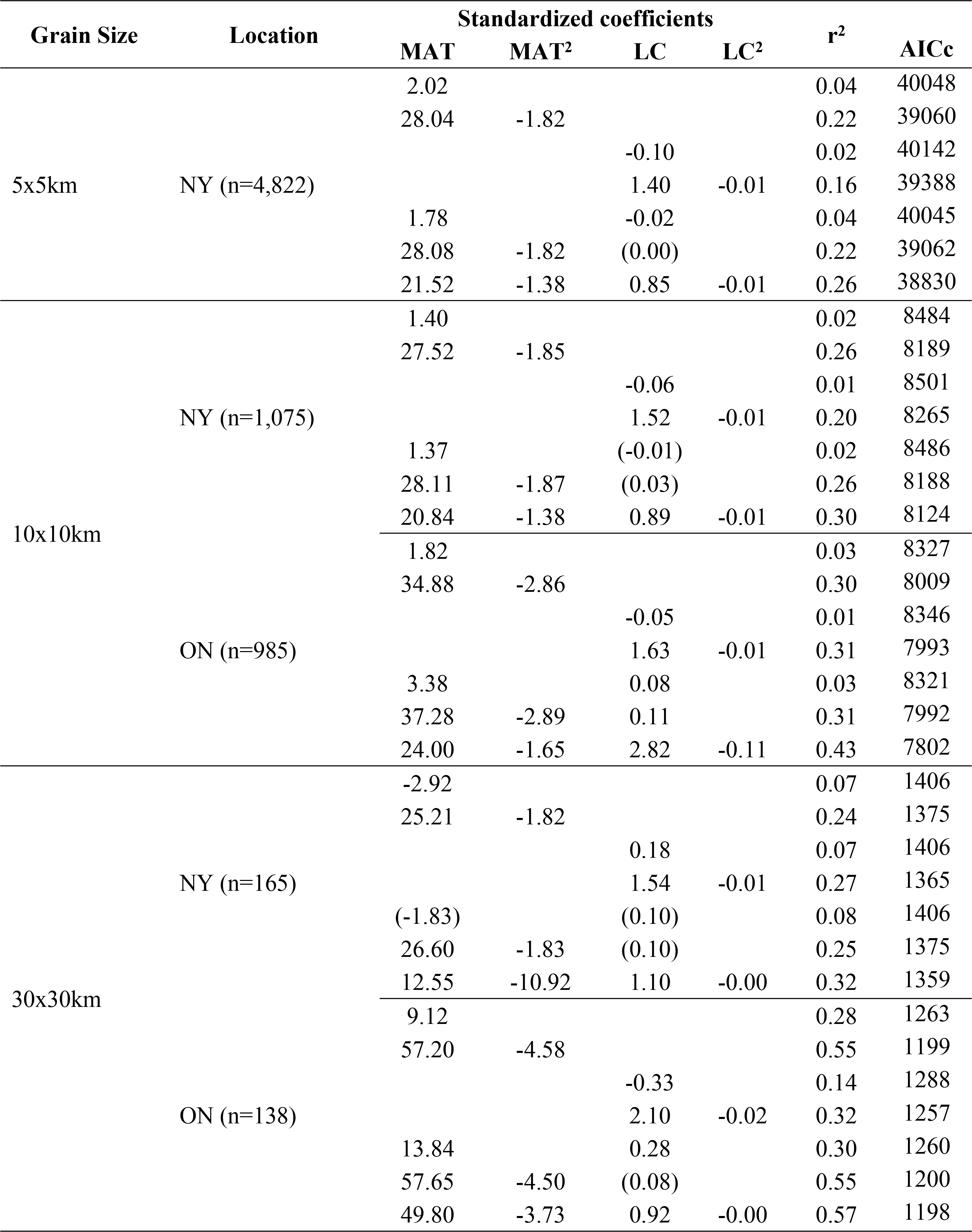
OLS regressions of **censused richness** as a function of temperature (MAT) and the proportion of natural land cover (LC) in grid cells in southern Ontario and New York State. Conventions as in Table 1.

**Fig 4.**
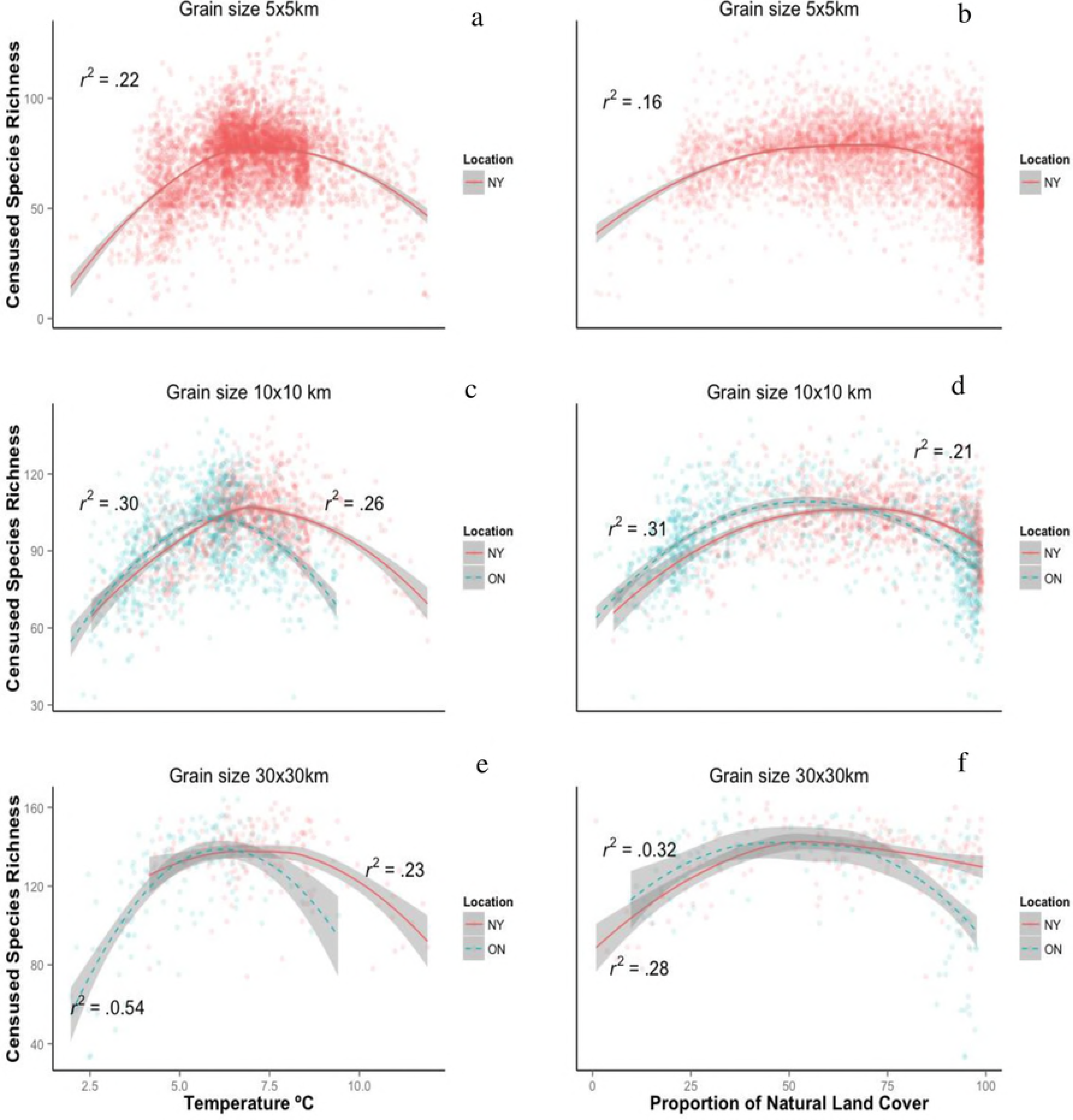
Distribution of censused avian species richness in a) 4,822 cells of 25-km^2^ in NY, b) 2,060 cells of 100-km^2^ in ON nad NY, and c) 303 cells of 900-km^2^ in ON and NY

**Fig 5.**
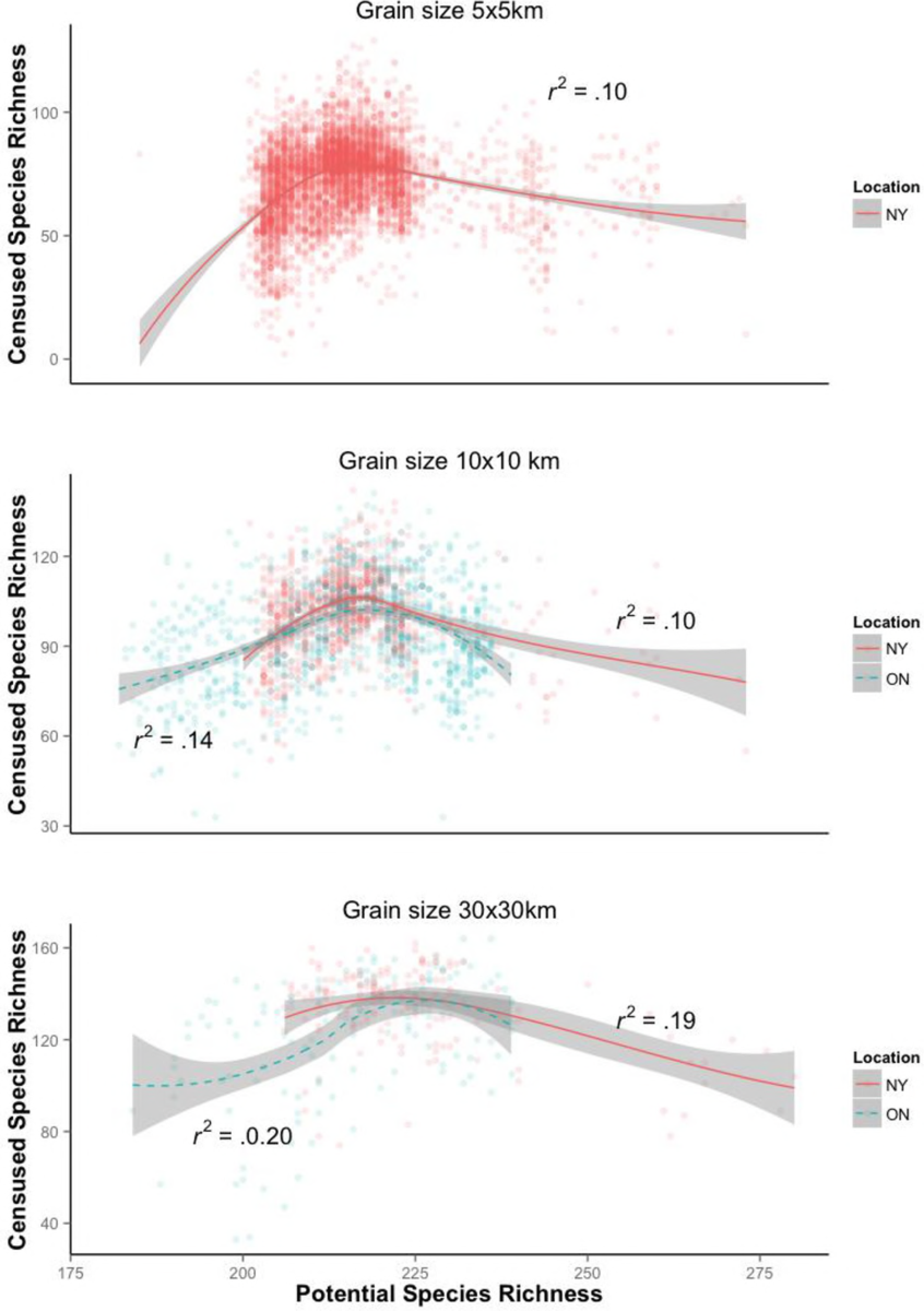
Censused species richness generated from atlases as a function of temperature (a,c,e) and the proportion of natural land cover (b,d,f) in grid cells covering southern Ontario and New York State at different spatial grain sizes. R2 represents the goodness of fit of OLS regression models. Richness peaks roughly at 53% (NY, 5×5km), 65% (NY, 10×10km), 55% (ON, 10×10km), 64% (NY, 30×30km), and 52% (ON, 30×30km) of natural land cover.

*Censused richness* is only weakly related to *range-map richness* over this study area (Fig 6), and the relationship is not monotonic, which is surprising since range-map richness in principle corresponds to the species pool available in the region [54].

**Fig 6.**
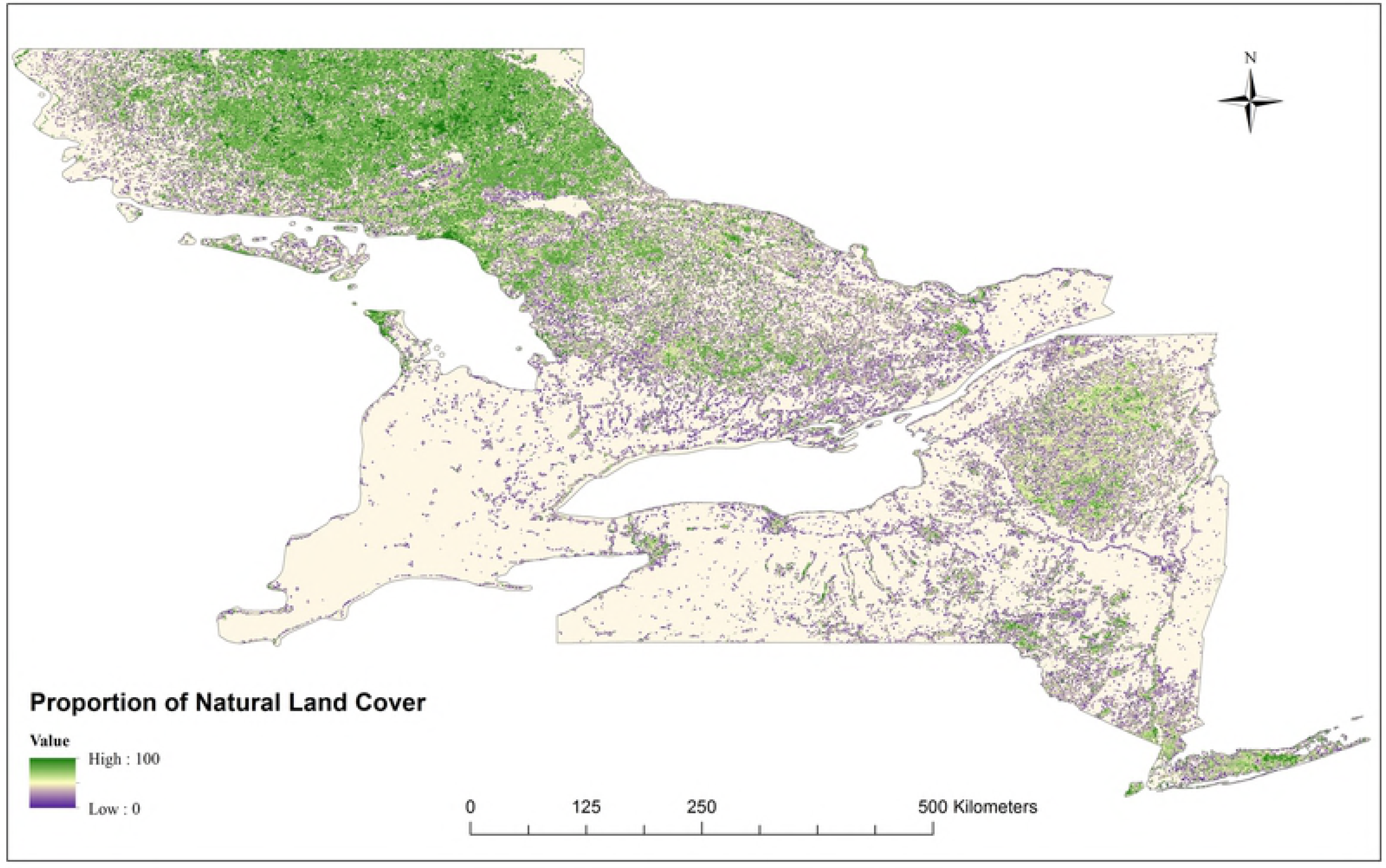
Censused richness as a function of range-map richness at different spatial grain sizes. a) n=4,822 in NY (5×5km), b) 985 and 1,075 for ON and NY, respectively, at 10×10km scale, and 251 covering ON and NY (30×30 km).

An effect of land cover on censused richness is most apparent in multiple regression models, after controlling for range-map richness (Table 3). Yet, the relationship is peaked with maximum censused richness reached roughly between 52-65% natural land cover, depending on the data type and grain size (Fig 5b,d,f). These results are only consistent with the proposition that protecting natural cover protects richness at very low natural cover.

**Table 3.**
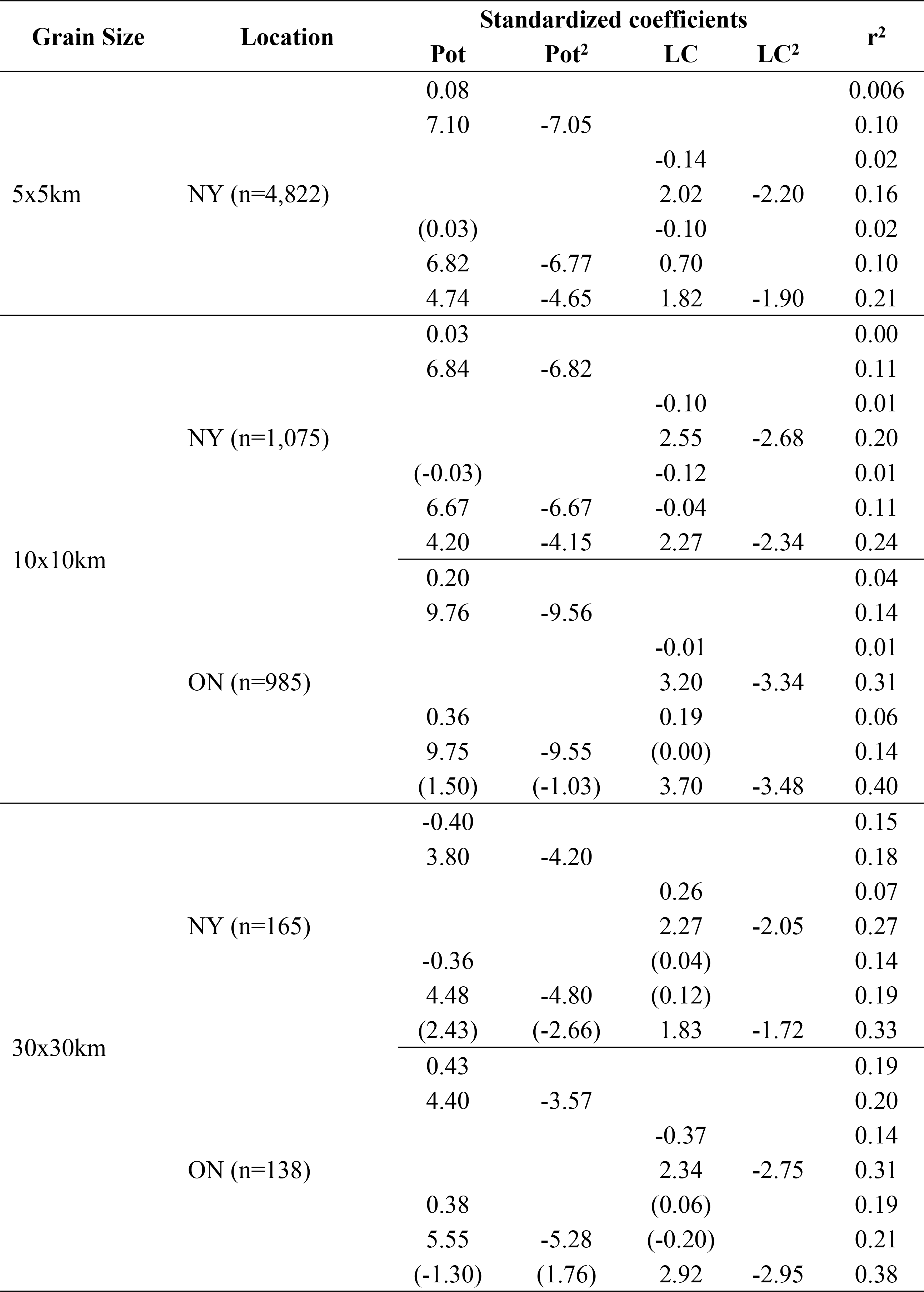
Multiple regressions between censused richness and land cover (LC), accounting for range-map richness (Pot).

Spatial autoregressive models do not change the qualitative patterns described above. Overall, these models performed better than the OLS models based on AIC comparisons (Tables 1, 2 main text vs. S1-S2 Tables). However, incorporating spatial autocorrelation increased variance explained of range-map richness models by ≤ 1% (comparison of r-squares from Table 1 vs. S1 Table), and the regression coefficients differ little. For censused richness, the additional variance explained added by autoregressive error models was also very small, but it seems to be more relevant at the 30×30km cell size (comparison of r-square from Table 2 vs. S2 Table).

## DISCUSSION

There is a substantial body of conservation literature that relates species’ distributions to land cover features (i.e., forest cover, fragmentation, land-use intensity, etc) at national or global scales in order to draw conclusions about the effect of habitat loss to biodiversity. The overall message from this type of research is that habitat loss is a major cause of species loss. For example, Betts et al. [45] tested the effects of deforestation on IUCN-red listed species. The authors found that deforestation greatly increases the likelihood of a species to be threatened worldwide, and predicted that up to roughly 220 species of vertebrates will fall within threatened categories in the next 30 years due to forest conversion in tropical regions. Similarly, Tilman et al. [55] reviewed the future threats to biodiversity and argued that between 40-80% of all threatened mammal and bird species are imperilled due to broad-scale habitat loss (i.e., agriculture, logging and development, their Figure 1a). Both studies used IUCN/Bird Life International coarse-grained range maps to arrive at their conclusions.

Yet, when examined closely, richness generated from range maps relates *negatively* to the proportional of natural land cover (i.e., mostly forest cover), probably because the patterns of richness are mainly driven by temperature (Fig 3, Table 1). Land cover varies dramatically over relatively fine spatial scales (Fig 1), but species’ ranges do not become peppered with areas of absence. Given that ranges circumscribe occupied and unoccupied areas, there is little reason to expect them to respond to land cover changes. Rather, we suspect that the correlations observed by Betts et al. [45] and by Tilman et al. [55] between range-map richness of <u>red-listed</u> species and natural land cover primarily reflect the fact that recent habitat loss within a species’ range is a criterion used by IUCN to define red-listed species (IUCN Standards and Petitions Subcommittee, 2017). The same issue can be observed in other assessments of threats to biodiversity that nearly always include human-induced destruction of natural habitats [1,55,57,58], introducing circularity into the proposed causal link between habitat loss and biodiversity loss.

In contrast, the pattern we found in range-map richness is consistent with other studies showing that, at coarse-grain and broad-extent, richness is strongly related (r^2^ ≅ 0.7-0.8) to current climatic variables [22,59,60]. While the mechanisms underlying such patterns are contentious [32,33,61,62], richness of most species groups increases monotonically with mean annual temperature (MAT) and/or precipitation [22]. Although the richness-climate relationship may vary somewhat among taxonomic groups [63], and among regions [31,64], climate always explain large amount of the variance and is congruent across space and time [61].

If the conversion of natural areas to human-dominated landscapes makes those areas completely unavailable to species, then there should be a positive spatial relationship between species richness and the total area of remnants of natural areas, irrespectively of spatial grain (e.g., [65]). Based on precisely this assumption, species-area relationships (SARs) have frequently been used to forecast species losses (e.g. number of species extinct or threatened) from removal of natural (usually forested) cover [27,39,42,44,66]. At coarse spatial grains and large extents, those forecasts have greatly exceeded observed species losses [7,27,65,67]. The discrepancy is sometimes attributed to “extinction debt”: extinctions that are predicted to occur, but that have not had time to do so. The difficulty is that the concept of “extinction debt” assumes that the causal link between species extinction and habitat loss exists, despite data to the contrary. Nonetheless, species-areas relationships are still commonly applied in conservation studies to predict loss of species as a function of habitat modification [7,27,42,66,68].

While it is generally accepted that richness is determined by climatic processes at coarser grains, environmental disturbances and stochastic processes may play a role in determining number of species at smaller grains [19]. Newbold et al. (2015) suggest that land use and land-use intensity may have major consequences for global biodiversity at local scales. They find that within high-disturbed sites (i.e., urban, pasture, croplands) richness can be reduced by >70%. It is not surprising that diversity is low in highly disturbed landscapes [10,69–72]. However, at the landscape scale, moderate habitat conversion appears *not* to be associated with species loss. For example, bird species richness peaks at intermediate amounts of natural land cover in southern Ontario, Canada [73], and at intermediate availability of trees in landscapes of Florence, Italy [74]. In our analysis, censused (Fig. 5b,d,f) species richness does not increase monotonically as a function of the proportion of natural habitat at any sampled grain size. Rather, censused richness-land cover relationships are peaked, leading to the conclusion that land cover is important to maintain biodiversity at very low natural cover, at its best. Our results are in striking contrast to a study by Betts et al. [45], which argues that the loss of biodiversity should be highest when natural cover is lost in fully forested areas.

What, then, causes the peaked relationship between avian richness and natural land cover? Using the same dataset as ours for southern Ontario, Desrochers et al. [73]proposed that the peaked relationship results from the sum of two SAR curves: the number of forest birds increases with the amount of forest cover, while the richness of open-habitat birds species increases with human-dominated areas. Thus, habitat heterogeneity may increase richness within landscapes [73]. De Camargo & Currie [75] further detailed the relationship by proposing a modified SAR models to predict avian species occurring in human-dominated landscapes in southern Ontario. Their model proposes there are some parts of the human-dominated land covers that are “available” for bird species to be present, while some other parts of the landscape are simply “lost”, and no species can thrive there. Because of the environmental similarities between Ontario and New York State, in term of biomes and climatic conditions, it is reasonable to assume that the same sort of mechanisms should be shaping the peaked richness-land cover relationships in New York State.

Habitat loss *per se* may not be the biggest threat to biodiversity. Instead, other factors may play a major role in explaining diversity decline, or act synergically with habitat loss to imperil species [76]. For instance, hunting practices have been linked to pre-historic [77] and modern [78] species extinctions. Yet hunting is poorly represented in assessments of threats to biodiversity [79]. Pesticides have been related to species losses in agricultural landscapes [6,80,81]. Also, land-use intensity has emerged as a potential main driver of species decline worldwide [2,23,82–84]. South-eastern Ontario, and western NY State, are heavily agricultural. Long Island has high human population density. It is in these areas that censused richness is lowest. Habitat destruction could be the first environmental disturbance happening at a locality, followed by other human-induced environmental stressors, but it may not be the primary cause of species loss since the other factors may act independently to eliminate species.

In conclusion, our study suggests that conserving natural land cover *per se* would not necessarily conserve species diversity. Remaining natural land cover in landscape is a poor, inconsistent predictor of species richness. Censused richness only seems to be negatively affected at very low levels natural land cover. Clearly, better environmental predictors of diversity changes are necessary, instead of aiming for “evidence complacency” [85] by creating a panchreston around habitat loss.

## ACKNOWLEDGMENTS

Thanks to the sponsors of the Atlas of the Breeding Bird of Ontario: Bird Studies Canada, Canadian Wildlife Service, Federation of Ontario Naturalists, Ontario Field Ornithologists, and Ontario Ministry of Natural Resources for supplying Atlas data, and to the thousands of volunteer participants who gathered data. The Natural Sciences and Engineering Research Council of Canada supported this work.

## SUPPORTING INFORMATION

S1 Fig. Linear regressions between the proportion of natural land cover and temperature at different spatial grain sizes.

S2 Fig. Censused richness-natural land cover relationships for the warmest and coldest places in southern Ontario and New York State. a) Coldest places in New York State (5×5km − n=1,097, 10×10km − n=244, 30×30km − n=28 cells); b) Warmest places in New York (5×5km − n=240, 10×10km − n=40, 30×30km − n=15 cells); c) Coldest places in Ontario (10×10km − n=243, 30×30km − n=46 cells); d) Warmest places in Ontario (10×10km − n=291, 30×30km − n=26 cells).

**S1 Table**. Statistical outcome of **spatial autocorrelation** models between potential richness as a function of temperature (MAT) and the proportion of natural land cover (LC) in grid cells of southern Ontario and New York State. * - term statistically not significant.

**S2 Table.** Statistical outcome **spatial autocorrelation** models between **realised richness** as a function of temperature (MAT) and the proportion of natural land cover (LC) in grid cells of southern Ontario and New York State. * - term statistically not significant.

S3 Fig. Potential species richness generated from range maps as a function of temperature (a,c,e) and the proportion of natural land cover (b,d,f) in grid cells covering southern Ontario and New York State at different spatial grain sizes. R^2^ represents the goodness of fit of second degree polynomial OLS regression models.

## REFERENCES

1. Wilcove DS, Rothstein D, Dubow J, Phillips A, Losos E. Quantifying threats to imperiled species in the United States. Bioscience. JSTOR; 1998;48: 607–615. Available: http://www.jstor.org/stable/1313420

2. Pekin BK, Pijanowski BC. Global land use intensity and the endangerment status of mammal species. Mac Nally R, editor. Divers Distrib. 2012;18: 909–918. doi:10.1111/j.1472-4642.2012.00928.x

3. Baillie J, Hilton-Taylor C, Stuart SN. A Global Species Assessment [Internet]. IUCN Publications Services Unit. World Conservation Union; 2004. Available: http://books.google.com/books?hl=en&lr=&id=Djr8v_-mFzYC&oi=fnd&pg=PR11&dq=2004+IUCN+Red+List+of+Threatened+Species+A+Global+Species+Assessment&ots=tmJjPmYsA9&sig=o1yHf2m6qgvKNdLYZ-RS5B5zvZE

4. Krauss J, Bommarco R, Guardiola M, Heikkinen RK, Helm A, Kuussaari M, et al. Habitat fragmentation causes immediate and time-delayed biodiversity loss at different trophic levels. Ecol Lett. 2010;13: 597–605. doi:10.1111/j.1461-0248.2010.01457.x

5. Sala OE, Chapin III S, Armesto JJ, Berlow E, Bloomfield J, Dirzo R, et al. Global Biodiversity Scenarios for the Year 2100. Science (80-). 2000;287: 1770–1774. doi:10.1126/science.287.5459.1770

6. Tilman D, Fargione J, Wolff B, D’Antonio C, Dobson a, Howarth R, et al. Forecasting agriculturally driven global environmental change. Science. 2001;292: 281–4. doi:10.1126/science.1057544

7. Pereira HM, Leadley PW, Proenca V, Alkemade R, Scharlemann JPW, Fernandez-Manjarres JF, et al. Scenarios for Global Biodiversity in the 21st Century. Science (80-). 2010;330: 1496–1501. doi:10.1126/science.1196624

8. Dooley JL, Bowers MA. Demographic Responses to Habitat Fragmentation: Experimental Tests at the Landscape and Patch Scale. Ecology. 1998;79: 969–980.

9. Prugh LR. An evaluation of patch connectivity measures. Ecol Appl. 2009;19: 1300–1310. doi:10.1890/08-1524.1

10. Guldemond R a. R, Van Aarde RJ. Forest patch size and isolation as drivers of bird species richness in Maputaland, Mozambique. J Biogeogr. 2010;37: 1884–1893. doi:10.1111/j.1365-2699.2010.02338.x

11. Martin AE, Fahrig L. Measuring and selecting scales of effect for landscape predictors in species-habitat models. Ecol Appl. 2012;22: 2277–2292. doi:10.1890/11-2224.1

12. Dale V, Brown S, Haeuber R, Hobbs N, Huntly N, Naiman R, et al. Ecological principles and guidelines for managing the use of land. Ecol Appl. 2000;10: 639–670. Available: http://www.esajournals.org/doi/abs/10.1890/1051-0761(2000)010%5B0639:EPAGFM%5D2.0.CO%3B2

13. Hurlbert AH, Jetz W. Species richness, hotspots, and the scale dependence of range maps in ecology and conservation. Proc Natl Acad Sci. 2007;104: 13384–13389. doi:10.1073/pnas.0704469104

14. Collinge SK. Effects of Grassland Fragmentation on Insect Species Loss, Colonization, and Movement Patterns. Ecology. 2000;81: 2211. doi:10.2307/177109

15. Husté A, Boulinier T. Determinants of local extinction and turnover rates in urban bird communities. Ecol Appl. 2007;17: 168–80. Available: http://www.ncbi.nlm.nih.gov/pubmed/17479843

16. D’Amen M, Bombi P. Global warming and biodiversity: Evidence of climate-linked amphibian declines in Italy. Biol Conserv. Elsevier Ltd; 2009;142: 3060–3067. doi:10.1016/j.biocon.2009.08.004

17. Arroyo-Rodríguez V, Dias PAD. Effects of habitat fragmentation and disturbance on howler monkeys: a review. Am J Primatol. 2010;72: 1–16. doi:10.1002/ajp.20753

18. Cousins S a. O, Vanhoenacker D. Detection of extinction debt depends on scale and specialisation. Biol Conserv. Elsevier Ltd; 2011;144: 782–787. doi:10.1016/j.biocon.2010.11.009

19. Ricklefs RE. A comprehensive framework for global patterns in biodiversity. Ecol Lett. 2004;7: 1–15. doi:10.1046/j.1461-0248.2003.00554.x

20. Newbold T. Ecological traits affect the response of tropical forest bird species to land-use intensity. Proc R Soc B Biol Sci. 2012;280: 2012–2131. Available: http://rspb.royalsocietypublishing.org/content/280/1750/20122131.short

21. Newbold T, Hudson LN, Phillips HRP, Hill SLL, Contu S, Lysenko I, et al. A global model of the response of tropical and sub-tropical forest biodiversity to anthropogenic pressures. Proc R Soc London Ser B Biol Sci. 2014;281: 20141371. doi:http://dx.doi.org/10.1098/rspb.2014.1371

22. Field R, Hawkins B a., Cornell H V., Currie DJ, Diniz-Filho JAF, Guégan J-F, et al. Spatial species-richness gradients across scales: a meta-analysis. J Biogeogr. 2009;36: 132–147. doi:10.1111/j.1365-2699.2008.01963.x

23. Newbold T, Hudson LN, Hill SLL, Contu S, Lysenko I, Senior RA, et al. Global effects of land use on local terrestrial biodiversity. Nature. 2015;520: 45–50. doi:10.1038/nature14324

24. Fahrig L. How much habitat is enough? Biol Conserv. 2001;100: 65–74. doi:10.1016/S0006-3207(00)00208-1

25. Estavillo C, Pardini R, Da Rocha PLB. Forest loss and the biodiversity threshold: An evaluation considering species habitat requirements and the use of matrix habitats. PLoS One. 2013;8: 1–10. doi:10.1371/journal.pone.0082369

26. Francesco Ficetola G, Denoël M. Ecological thresholds: an assessment of methods to identify abrupt changes in species-habitat relationships. Ecography (Cop). 2009;32: 1075–1084. doi:10.1111/j.1600-0587.2009.05571.x

27. Stork NE. Re-assessing current extinction rates. Biodivers Conserv. 2009;19: 357–371. doi:10.1007/s10531-009-9761-9

28. Algar AC, Kerr JT, Currie DJ. Evolutionary constraints on regional faunas: whom, but not how many. Ecol Lett. 2008; 57–65. doi:10.1111/j.1461-0248.2008.01260.x

29. Hawkins BA, Field R, Cornell HV, Currie DJ, Guégan JF, Kaufman DM, et al. Energy, water, and broad-scale geographic patterns of species richness. Ecology. Eco Soc America; 2003;84: 3105–3117. doi:10.1890/03-8006

30. Currie DJ. Energy and Large Scale Patterns of Animal and Plant Species Richness. Am Nat. JSTOR; 1991;137: 27–49. Available: http://www.jstor.org/stable/2462155

31. Francis AP, Currie DJ. A globally consistent richness-climate relationship for angiosperms. Am Nat. UChicago Press; 2003;161: 523–536. Available: http://www.journals.uchicago.edu/doi/pdf/10.1086/368223

32. Romdal TS, Araújo MB, Rahbek C. Life on a tropical planet: niche conservatism and the global diversity gradient. Glob Ecol Biogeogr. 2013;22: 344–350. doi:10.1111/j.1466-8238.2012.00786.x

33. Wiens JJ, Ackerly DD, Allen AP, Anacker BL, Buckley LB, Cornell H V., et al. Niche conservatism as an emerging principle in ecology and conservation biology. Ecol Lett. 2010;13: 1310–1324. doi:10.1111/j.1461-0248.2010.01515.x

34. Findlay CS, Houlahan J. Anthropogenic Correlates of Species Richness in Southeastern Ontario Wetlands. Conserv Biol. 1997;11: 1000–1009. doi:10.1046/j.1523-1739.1997.96144.x

35. Smith AC, Fahrig L, Francis CM. Landscape size affects the relative importance of habitat amount, habitat fragmentation, and matrix quality on forest birds. Ecography (Cop). 2011;34: 103–113. doi:10.1111/j.1600-0587.2010.06201.x

36. Schmiegelow FKA, Mönkkönen M. Habitat loss and fragmentation in dynamic landscape: avian perspectives from the boreal forest. Ecol Appl. 2002;12: 375–389. doi:10.1890/1051-0761(2002)012[0375:HLAFID]2.0.CO;2

37. Drapeau P, Leduc A, Giroux J. Landcape-Scale Disturbances and Changes in Bird Communities of Boreal Mixed-Forest Forests. Ecol Monogr. 2000;70: 423–444. Available: http://www.esajournals.org/doi/abs/10.1890/0012-9615(2000)070%5B0423:LSDACI%5D2.0.CO%3B2

38. Tilman D, May RM, Lehman CL, Nowak MA. Habitat destruction and the extinction debt. Nature. Nature Publishing Group; 1994;371: 65–66. doi:10.1038/371065a0

39. Wilson EO. The current state of biological diversity. Biodiversity. 1988. pp. 3–18.

40. Pimm SL, Raven P. Extinction by numbers. Nature. 2000;403: 843–845.

41. Brooks TM, Mittermeier R a., Mittermeier CG, da Fonseca G a. B, Rylands AB, Konstant WR, et al. Habitat Loss and Extinction in the Hotspots of Biodiversity. Conserv Biol. 2002;16: 909–923. doi:10.1046/j.1523-1739.2002.00530.x

42. Pimm S, Raven P, Peterson A, Sekercioglu CH, Ehrlich PR. Human impacts on the rates of recent, present, and future bird extinctions. Proc Natl Acad Sci U S A. 2006;103: 10941–6. doi:10.1073/pnas.0604181103

43. Pimm SL, Jenkins CN, Abell R, Brooks TM, Gittleman JL, Joppa LN, et al. The biodiversity of species and their rates of extinction, distribution, and protection. Science. 2014;344: 1246752. doi:10.1126/science.1246752

44. Hubbell SP, He F, Condit R, Borda-de-Agua L, Kellner J, ter Steege H. How many tree species are there in the Amazon and how many of them will go extinct? Proc Natl Acad Sci. 2008;105: 11498–11504. doi:10.1073/pnas.0801915105

45. Betts MG, Wolf C, Ripple WJ, Phalan B, Millers KA, Duarte A, et al. Global forest loss disproportionately erodes biodiversity in intact landscapes. Nature. Nature Publishing Group; 2017; doi:10.1038/nature23285

46. Cadman MD, Sutherland DA, Beck GG, Lepage D, Couturier AR. Atlas of the Breeding Birds of Ontario, 2001-2005. The Auk. 2009. pp. 469–472. doi:10.1525/auk.2009.4409.3

47. McGowan KJ, Corwin K. The Second Atlas of Breeding Birds in New York State. First Edit. McGowan KJ, Corwin K, editors. Cornell University Press; 2008.

48. Tuanmu MN, Jetz W. A global 1-km consensus land-cover product for biodiversity and ecosystem modelling. Glob Ecol Biogeogr. 2014;23: 1031–1045. doi:10.1111/geb.12182

49. Trzcinski M, Fahrig L, Merriam G. Independent Effects of Forest Cover and Fragmentation on the Distribution of Forest Breeding Birds. Ecol Appl. 1999;9: 586–593. Available: http://www.jstor.org/stable/2641146

50. Fick SE, Hijmans RJ. WorldClim 2: New 1-km spatial resolution climate surfaces for global land areas. Int J Climatol. 2017; doi:10.1002/joc.5086

51. R Development Core Team R. R: A Language and Environment for Statistical Computing. R Foundation for Statistical Computing. 2011. doi:10.1007/978-3-540-74686-7

52. Diniz-Filho JAF, Bini LM, Hawkins B a. Spatial autocorrelation and red herrings in geographical ecology. Glob Ecol Biogeogr. 2003;12: 53–64. doi:10.1046/j.1466-822X.2003.00322.x

53. Kissling WD, Carl G. Spatial autocorrelation and the selection of simultaneous autoregressive models. Glob Ecol Biogeogr. 2007;17: 57–71. doi:10.1111/j.1466-8238.2007.00334.x

54. Cornell H V, Harrison SP. What Are Species Pools and When Are They Important? Annu Rev Ecol Evol/ Syst. 2014;45: 45–67. doi:10.1146/annurev-ecolsys-120213-091759

55. Tilman D, Clark M, Williams DR, Kimmel K, Polasky S, Packer C. Future threats to biodiversity and pathways to their prevention. Nature. 2017;546: 73–81. doi:10.1038/nature22900

56. IUCN Standards and Petitions Subcommittee. Guidelines for using the IUCN Red List categories and criteria. Version 11. Retrieved from http://www.iucnredlist.org/documents/RedListGuidelines.pdf Last accessed 27 March 2017. 2017;

57. Czech B, Krausman PR, Devers PK. Economic associations among causes of species endangerment in the United States. Bioscience. Univ California Press; 2000;50: 593–601. doi:10.1641/0006-3568(2000)050

58. Venter O, Brodeur NN, Nemiroff L, Belland B, Dolinsek IJ, Grant JW a. Threats to Endangered Species in Canada. Bioscience. 2006;56: 903. doi:10.1641/0006-3568(2006)56[903:TTESIC]2.0.CO;2

59. Hillebrand H. On the Generality of the Latitudinal Diversity Gradient. Am Nat. 2004;163: 192–211. doi:10.1086/381004

60. Buckley LB, Hurlbert AH, Jetz W. Broad-scale ecological implications of ectothermy and endothermy in changing environments. Glob Ecol Biogeogr. 2012;21: 873–885. doi:10.1111/j.1466-8238.2011.00737.x

61. Boucher-Lalonde V, De Camargo RX, Fortin J-M, Khair S, So RI, Vázquez Rivera H, et al. The weakness of evidence supporting tropical niche conservatism as a main driver of current richness-temperature gradients. Glob Ecol Biogeogr. 2015;24: 795–803. doi:10.1111/geb.12312

62. Mittelbach GG, Schemske DW, Cornell H V, Allen AP, Brown JM, Bush MB, et al. Evolution and the latitudinal diversity gradient: speciation, extinction and biogeography. Ecol Lett. 2007;10: 315–31. doi:10.1111/j.1461-0248.2007.01020.x

63. Wolters V, Wolters V, Bengtsson J, Bengtsson J, Zaitsev AS. Relationship Among The Species Richness Of Diffenrent Taxa. Ecology. 2006;87: 1886–1895. doi:10.1890/0012-9658(2006)87[1886:RATSRO]2.0.CO;2

64. Jiménez I, Ricklefs RE. Diversity anomalies and spatial climate heterogeneity. Glob Ecol Biogeogr. 2014;23: 988–999. doi:10.1111/geb.12181

65. He F, Hubbell SP. Species-area relationships always overestimate extinction rates from habitat loss. Nature. 2011;473: 368–371. doi:10.1038/nature09985

66. Pimm SL, Jenkins CN, Abell R, Brooks TM, Gittleman JL, Joppa LN, et al. The biodiversity of species and their rates of extinction, distribution, and protection. Science. 2014;344: 1246752. doi:10.1126/science.1246752

67. Pereira HM, Borda-de-Água L, Martins IS. Geometry and scale in species-area relationships. Nature. 2012;482. doi:10.1038/nature10857

68. Haddad NM, Gonzalez A, Brudvig LA, Burt MA, Levey DJ, Damschen EI. Experimental evidence does not support the Habitat Amount Hypothesis. Ecography (Cop). 2017;125: 336–342. doi:10.1111/ecog.02535

69. Kajzer J, Lenda M, Kośmicki A, Bobrek R, Kowalczyk T, Martyka R, et al. Patch occupancy and abundance of local populations in landscapes differing in degree of habitat fragmentation: A case study of the colonial black-headed gull, Chroicocephalus ridibundus. J Biogeogr. 2012;39: 371–381. doi:10.1111/j.1365-2699.2011.02604.x

70. Norris D, Michalski F, Peres C a. Habitat patch size modulates terrestrial mammal activity patterns in Amazonian forest fragments. J Mammal. 2010;91: 551–560. doi:10.1644/09-MAMM-A-199.1

71. Fahrig L. Rethinking patch size and isolation effects: The habitat amount hypothesis. J Biogeogr. 2013;40: 1649–1663. doi:10.1111/jbi.12130

72. Bender DJ, Contreras TA, Fahrig L. Habitat loss and population decline: a meta-analysis of the patch size effect. Ecology. Eco Soc America; 1998;79: 517–533. doi:10.1890/0012-9658(1998)079[0517:HLAPDA]2.0.CO;2

73. Desrochers RE, Kerr JT, Currie DJ. How, and how much, natural cover loss increases species richness. Glob Ecol Biogeogr. 2011;20: 857–867. doi:10.1111/j.1466-8238.2011.00658.x

74. Chiari C, Dinetti M, Licciardello C, Licitra G, Pautasso M. Urbanization and the more-individuals hypothesis. J Anim Ecol. 2010;79: 366–371. doi:10.1111/j.1365-2656.2009.01631.x

75. De Camargo RX, Currie DJ. An empirical investigation of why species - area relationships overestimate species losses. Ecology. 2015;96: 1253–1263.

76. Brook BW, Sodhi NS, Bradshaw CJ a. Synergies among extinction drivers under global change. Trends Ecol Evol (Personal Ed. 2008;23: 453–60. doi:10.1016/j.tree.2008.03.011

77. Malhi Y, Doughty CE, Galetti M, Smith FA, Svenning J-C, Terborgh JW. Megafauna and ecosystem function from the Pleistocene to the Anthropocene. Proc Natl Acad Sci. 2016;113: 838–846. doi:10.1073/pnas.1502540113

78. Corlett RT. The impact of hunting on the mammalian fauna of tropical Asian forests. Biotropica. 2007;39: 292–303. doi:10.1111/j.1744-7429.2007.00271.x

79. Joppa LN, O’Connor B, Visconti P, Smith C, Geldmann J, Hoffmann M, et al. Filling in biodiversity threat gaps. Science (80-). 2016;352: 416–418. doi:10.1126/science.aaf3565

80. Gibbs KE, Mackey RL, Currie DJ. Human land use, agriculture, pesticides and losses of imperiled species. Divers Distrib. John Wiley & Sons; 2009;15: 242–253. doi:10.1111/j.1472-4642.2008.00543.x

81. Coristine LE, Kerr JT. Habitat loss, climate change, and emerging conservation challenges in Canada. 2011;451: 435–451. doi:10.1139/Z11-023

82. Tylianakis JM, Klein AM, Lozada T, Tscharntke T. Spatial scale of observation affects α, β and γ diversity of cavity-nesting bees and wasps across a tropical land-use gradient. J Biogeogr. 2006;33: 1295–1304. doi:10.1111/j.1365-2699.2006.01493.x

83. Hendrickx F, Maelfait J-P, Van Wingerden W, Schweiger O, Speelmans M, Aviron S, et al. How landscape structure, land-use intensity and habitat diversity affect components of total arthropod diversity in agricultural landscapes. J Appl Ecol. 2007;44: 340–351. doi:10.1111/j.1365-2664.2006.01270.x

84. Kehoe L, Senf C, Meyer C, Gerstner K, Kreft H, Kuemmerle T. Agriculture rivals biomes in predicting global species richness. Ecography (Cop). 2016; 1118–1128. doi:10.1111/ecog.02508

85. Sutherland WJ, Wordley CFR. Evidence complacency hampers conservation. Nat Ecol Evol. Springer US; 2017;1: 1215–1216. doi:10.1038/s41559-017-0244-1

